# Factors affecting microbial community composition in Split Lake, MB, Canada

**DOI:** 10.64898/2026.11.02.763575

**Authors:** Thamali Vidanage, Tataskweyak Environmental Monitoring Agency, R. Eric Collins

## Abstract

Split Lake is a large freshwater lake in northern Manitoba, Canada, that receives major inflows from the Nelson River, Burntwood River, and Grassy River. In 1976, during the Churchill River Diversion (CRD) project, most of the water in the Churchill River was diverted into the Nelson River via the Rat River, and the Burntwood River, creating a large inflow pressure in the lower Nelson area, including Split Lake. Decades after its implementation, the Churchill River Diversion still influences ecological and environmental processes in the area. In this study, metabarcoding of the 16S rRNA gene was employed to characterize the microbial community structure of the lake. Environmental factors (temperature, pH, turbidity, dissolved oxygen) and cyanotoxin levels were also assessed.

The objectives of this study were to establish a baseline for microbiota across the lake and examine how environmental variables affect microbial population dynamics. The study was conducted in the summer of 2024, encompassing 14 sites on Split Lake and 2 sites on Assean Lake, an adjacent waterbody with minimal influence from hydroelectric infrastructure. Split Lake was found to harbor six distinct microbial clusters, with turbidity and dissolved oxygen levels correlating with different aquatic regimes within the lake. The predominant taxa in Split Lake included Actinobacteria hgcI clade and *Cyanobium*. Microcystin concentrations measured throughout the study were always below the Maximum Acceptable Concentration (MAC) set for Canadian federal drinking water standards.

## 1 Introduction

Split Lake is a shallow, seasonally ice-covered riverine lake that spans approximately 46 km of the Nelson River in northern Manitoba, Canada. Tataskweyak is a remote community of about 2,300 residents located on the north shore of Split Lake. In May 2017, a boil water advisory was issued for the Tataskweyak Cree Public Water System (#6602), becoming a long-term advisory in May 2018 (Indigenous Services Canada 2025). Since 1978, Manitoba Hydro has operated the Churchill River Diversion (CRD) water management project, which redirects 75% of the Churchill River drainage system into the Burntwood River and then into the Nelson River at Split Lake (Duncan and Williamson, 1988). The CRD diversion increased flow in the Nelson River by approximately 32% (Wiens, 1994), resulting in extensive shoreline erosion of clay and silt sediments (Manitoba Hydro, 2015). In response to these impacts, researchers from the University of Manitoba and Land Guardians from the Tataskweyak Environmental Monitoring Agency co-developed this Community-Led Monitoring project with the goal of understanding changes in water quality observed by community members and to monitor for future harms to the aquatic ecosystems. The specific objectives of this study were to establish a baseline for microbial community structure in Split Lake, identify environmental drivers of microbial community structure, and explore harmful algal bloom risks in the lake.

Freshwater microorganisms play a crucial role in biogeochemical cycles and influence water quality in lakes (Newton et al., 2011). They also act as primary sources of food and nutrients for higher trophic level organisms (Islam et al., 2019). Their absence distorts the food chain, disrupting ecosystem functioning and ultimately affecting the balance of biotic and abiotic systems in lakes (Joshi, 2016). Microorganisms in aquatic environments are highly sensitive to environmental changes and human-induced stressors (Islam et al., 2019). Anthropogenic activities such as agriculture, fishing, and tourism affect the structure and function of the microbial community, which leads to pathogenic bacteria being introduced into the lakes (Islam et al., 2019). Bacteria are more sensitive to anthropogenic changes than fungi (Luo et al., 2022) and are therefore widely used as indicators in assessments of water quality worldwide (Dufour, 1984; Gronewold et al., 2008; Méndez et al., 2004; Wen et al., 2020). Freshwater bacterial community data provide valuable insights into the effects of land use on stream ecosystems and can serve as a complementary tool for screening stream catchment attributes (Hermans et al., 2024). Human activities have profoundly altered natural river connectivity by disrupting longitudinal and lateral linkages through structures such as dams and levees, and by indirectly modifying hydrological, thermal, and sediment regimes that shape riverine ecosystem processes (Nilsson and Berggren, 2000). Damming is the most significant human-induced disturbance to rivers, with about 70% of the world’s rivers being dammed (Grill et al., 2019). Studies in dam-impacted reservoirs have documented pronounced longitudinal shifts in bacterioplankton community structure, which are ultimately linked to increased water residence time within impoundments (Mašín et al., 2003).

Freshwater cyanobacterial harmful algal blooms represent a significant risk to human health and aquatic ecosystems, with evidence indicating that their occurrence and intensity are increasing worldwide (Bramburger et al., 2023). Nowicka-Krawczyk et al. (2022) showed that the emergence of cyanobacterial harmful algal blooms (CyanoHAB) is linked with the unsustainable use of water bodies and land by humans. Such changes in aquatic communities can further drive eutrophication processes by affecting trophic interactions, changing nutrient ratios, and promoting the movement of nutrients from benthic layers to the overlying water column (Schindler, 2006). In addition, CyanoHABs pose serious socioeconomic and ecological costs that impact the water quality, habitats, food webs, and tourism.

These bloom-forming genera can produce cyanotoxins (microcystin, cylindrospermopsin, nodularin) that can cause acute or chronic health effects for mammals and other organisms (Carmichael and Boyer, 2016; Metcalf and Codd, 2012). Cyanobacteria are widely recognized as key indicators of eutrophication in freshwater systems, as their blooms frequently occur in waters affected by excessive nutrient loading (Thajuddin and Dhanasekaran, 2016). As such, human-driven changes in water physicochemical properties can lead to localized sorting of aquatic bacterial communities (Chen et al., 2018), which in turn results in significant spatiotemporal variability in bacteria-mediated processes within aquatic ecosystems (Islam et al., 2019).

Using bacterial 16S rRNA gene metabarcoding, this study aimed to (1) establish a baseline characterization of the microbial community structure in the surface waters of Split Lake, northern Manitoba, a freshwater system influenced by hydroelectric development; (2) examine spatial variability in microbial community composition across the lake; (3) evaluate relationships between microbial community patterns and key physicochemical water characteristics; and (4) assess the potential of microbial metabarcoding data as an indicator of ecosystem disturbance associated with hydroelectric regulation.

## 2 Materials and methods

### 2.1 Site description and sampling

Surface water samples were collected by a small boat from 14 sites across Split Lake (56.2°N, 96.1°W) and 2 sites from Assean Lake (56.2°N, 96.5°W) from August 13–16, 2024 (Figure 1). At each site, approximately 1 L of water was filtered through 0.22 *µ*m Sterivex filters and stored frozen until DNA extraction. Sample temperature and pH were determined using a digital pH meter, and dissolved oxygen (DO) was determined using the CHEMets Dissolved Oxygen visual water testing kit. Turbidity was measured using the Palintest® Turbimeter Plus instrument. Secchi disc depth measurements were also taken at each site to assess water clarity.

**Figure 1.**
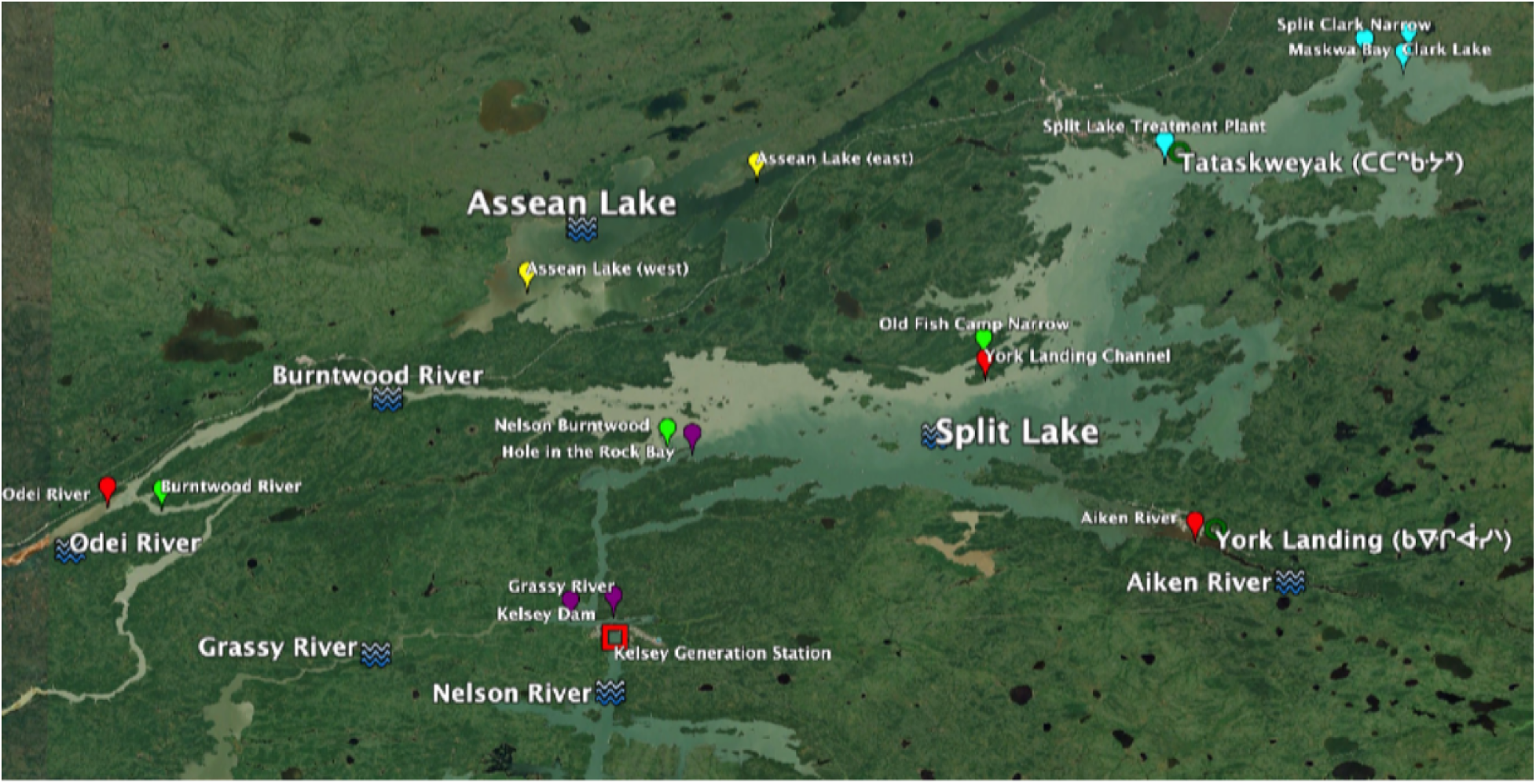
True color satellite image of the Split Lake region, Manitoba, taken on August 5, 2024, by the Sentinel-2 mission. Suspended sedimentary material (clay and silt) discolors the water, producing lighter shades. Water samples were collected by a small boat from August 13–16, 2024; pins are colored by hierarchical clustering of microbial community composition. Local rivers (blue waves) and Manitoba Hydro structures (red squares) are shown for context.

### 2.2 DNA extraction, amplicon library preparation, and processing

Qiagen DNeasy PowerSoil kits were used to extract DNA from Sterivex filters following the manufacturer’s protocols. Extracted DNA was quantified using a Qubit fluorometer and stored at *−*80 °C until sequencing. The V4 hypervariable region of the 16S rRNA gene was amplified using the 515F (GTGYCAGCMGCCGCG-GTAA) and 806RB (GGACTACNVGGGTWTCTAAT) primer pair. Amplicon libraries were sequenced using paired-end (2 *×* 250 bp) Illumina sequencing by CD Genomics (New York, USA). A total of 40k high-quality reads were obtained with a sequence depth range of 29,686 to 73,860 reads. Raw sequencing reads were quality-controlled by removing adapters and primer sequences using Cutadapt (Martin, 2011). Trimmed reads were then processed using the DADA2 (Callahan et al., 2016) package in R to infer amplicon sequence variants (ASVs), and taxonomic assignment was performed using the SILVA v138.2 reference database (Quast et al., 2013). Code for the bioinformatics pipeline is available at https://github.com/rec3141/microscape.

### 2.3 Statistical analysis

Statistical analyses and visualizations were performed in R. To explore the correlations between microbial community composition and environmental variables, Spearman correlation analysis was performed. The corrplot and pheatmap packages in R were used to produce heatmaps and correlation analysis visualizations, with p-values adjusted for multiple comparisons using the Benjamini–Hochberg method. Redundancy analysis (RDA) was performed to explore relationships between bacterial community composition and environmental factors using the vegan package (Oksanen et al., 2025).

### 2.4 Water chemistry analysis

The method American Society for Testing and Materials D6919-03 (ASTM, 2021) was followed for major cations (Mg^2+^, Ca^2+^, Na^+^), and EPA Method 300.0 (Pfaff, 1996) was used for anions (F*^−^*, Cl*^−^*, SO_4_^2*−*^). Concentrations were quantified using the Dionex 5000+ ion chromatography instrument at the Centre for Earth Observation Science, UCTEL, University of Manitoba. Standard reagents and consumables were sourced from Thermo Fisher Scientific.

### 2.5 Cyanotoxin analysis

The ABRAXIS® Microcystins Strip Test for Source Drinking Water was used to qualitatively screen microcystins in lake water samples following the manufacturer’s instructions. The sensitivity of the test kit is 1 ng/*µ*L or higher. All samples and reagents were allowed to reach room temperature before testing.

## 3 Results

### 3.1 Environmental parameters

Water temperature of the 16 sampling sites ranged between 20.6 °C and 23.2 °C (Table 1). Measurements of pH ranged from 7.2 to 8.0 and fell within federal guidelines (pH 7.0–8.7) for freshwater. The minimum federal guideline for dissolved oxygen in warm freshwater ecosystems is above 5.5 mg/L (Canadian Council of Ministers of the Environment 1999), which was met by all except one site (Maskwa Bay). The highest turbidity was observed in Burntwood River (65.00 NTU), while the lowest was observed in Assean Lake (west) (2.51 NTU).

**Table 1.** Metadata of sampling sites in Split Lake during the August 13^th^–16^th^, 2024 sampling expedition.

| Date | Site Name | Site ID | Longitude (deg E) | Latitude (deg W) | Temperature (°C) | pH | Dissolved Oxygen (ppm) | Turbidity (NTU) | Secchi depth (m) | Cluster color |
|---|---|---|---|---|---|---|---|---|---|---|
| 13/08/2024 | York Landing Channel | YLC | −96.2434 | 56.1472 | 23.2 | 8.0 | 8.5 | 5.78 | 1.50 | Red |
| 15/08/2024 | Aiken River | AK | −96.0824 | 56.0764 | 23.1 | 7.2 | 7.0 | 9.09 | 0.98 |  |
| 14/08/2024 | Odei River | OR | −96.9186 | 56.0905 | 21.1 | 7.8 | 7.5 | 34.70 | 0.47 |  |
| 16/08/2024 | Assean Lake (east) | ALe | −96.5961 | 56.1851 | 20.8 | 7.2 | 7.0 | 4.57 | 2.20 | Yellow |
| 16/08/2024 | Assean Lake (west) | ALw | −96.4192 | 56.2330 | 21.6 | 7.5 | 5.5 | 2.51 | 2.97 |  |
| 13/08/2024 | Nelson Burntwood | NB | −96.4879 | 56.1171 | 20.1 | 7.5 | 8.5 | 52.90 | 0.30 | Green |
| 14/08/2024 | Burntwood River | BR | −96.8768 | 56.0895 | 20.8 | 7.8 | 8.0 | 65.00 | 0.33 |  |
| 14/08/2024 | Old Fish Camp Narrow | OFCN | −96.2446 | 56.1565 | 21.6 | 7.2 | 8.0 | 40.10 | 0.39 |  |
| 16/08/2024 | Split Clark Narrow | SCN | −95.9498 | 56.2848 | 21.3 | 7.8 | 6.0 | 14.50 | 0.90 | Cyan |
| 16/08/2024 | Clark Lake | CL | −95.9198 | 56.2799 | 21.2 | 7.5 | 8.0 | 12.70 | 1.10 |  |
| 13/08/2024 | Split Lake Treatment Plant | SLTP | −96.1045 | 56.2409 | 21.2 | 7.5 | 10.0 | 19.10 | NA |  |
| 16/08/2024 | Maskwa Bay | MB | −95.9155 | 56.2890 | 20.8 | 7.5 | 5.0 | 13.60 | 1.08 |  |
| 13/08/2024 | Kelsey Dam | KD | −96.5290 | 56.0442 | 21.0 | 7.6 | 6.0 | 7.25 | 1.65 | Purple |
| 15/08/2024 | Hole in the Rock Bay | HRB | −96.4686 | 56.1148 | 21.1 | 7.5 | 6.0 | 7.94 | 1.58 |  |
| 13/08/2024 | Grassy River | GR | −96.5618 | 56.0424 | 20.6 | 7.8 | 6.0 | 6.47 | 1.70 |  |
| 15/08/2024 | Josey Spence Island | JSI | NA | NA | 21.6 | 7.5 | 7.0 | 36.60 | 0.42 | Magenta |

### 3.2 Water chemistry in the lake

Concentrations of major cations (Mg^2+^, Ca^2+^, Na^+^, K^+^) and anions (F*^−^*, Cl*^−^*, NO_3_*^−^*, SO_4_^2*−*^) were determined from each water sample (Table S1). Phosphates were not detected at any of the sampling sites. Sulfate concentrations were relatively high at Kelsey Dam (33.32 ppm) and Hole in the Rock Bay (28.66 ppm) sites, both located along the upper Nelson River watershed. Comparable sulfate levels were also observed at the York Landing Channel (28.65 ppm). In contrast, the lowest sulfate level was observed at Josey Spence Island (0.44 ppm); however, all measured ion concentrations remained below applicable federal guideline limits (Canadian Council of Ministers of the Environment, 2007).

### 3.3 Cyanotoxin concentration analysis

Microcystin concentrations measured across the sampling sites were consistently low. At York Landing Channel, Old Fish Camp Narrow, Nelson Burntwood, Kelsey Dam, Burntwood River, Assean Lake (west), Clark Lake, Hole in the Rock Bay, and Maskwa Bay, microcystin was not detected (0 *µ*g/L). Several sites recorded concentrations of 0.5 *µ*g/L, including Odei River, Assean Lake (east), Josey Spence Island, Split Clark Narrow, Split Lake Treatment Plant, Grassy River, and Aiken River. Importantly, none of the sites exceeded the maximum acceptable concentration established in the Canadian Water Quality Guidelines for drinking water.

### 3.4 Taxonomic abundance profiles

All quality-controlled sequences across 16 samples were resolved into 842 ASVs. Bray–Curtis dissimilarities among samples were calculated from fourth root transformed relative abundances and hierarchically clustered using the Ward.D algorithm in R using stats::hclust().

The sample dendrogram was cut into six clusters containing distinct community assemblages, including in waters strongly influenced by the Burntwood River watershed (green), upper Nelson River (purple), and Assean Lake (yellow), while samples from waters mostly influenced by the Odei River and Aiken River clustered together (red). Waters from the lower Nelson River watershed region of Split Lake, where inflowing river waters become well-mixed, were clustered with each other (cyan). Another site, “Josey Spence Island” (location information missing), clustered separately from all other samples (magenta; Figure 2).

**Figure 2.**
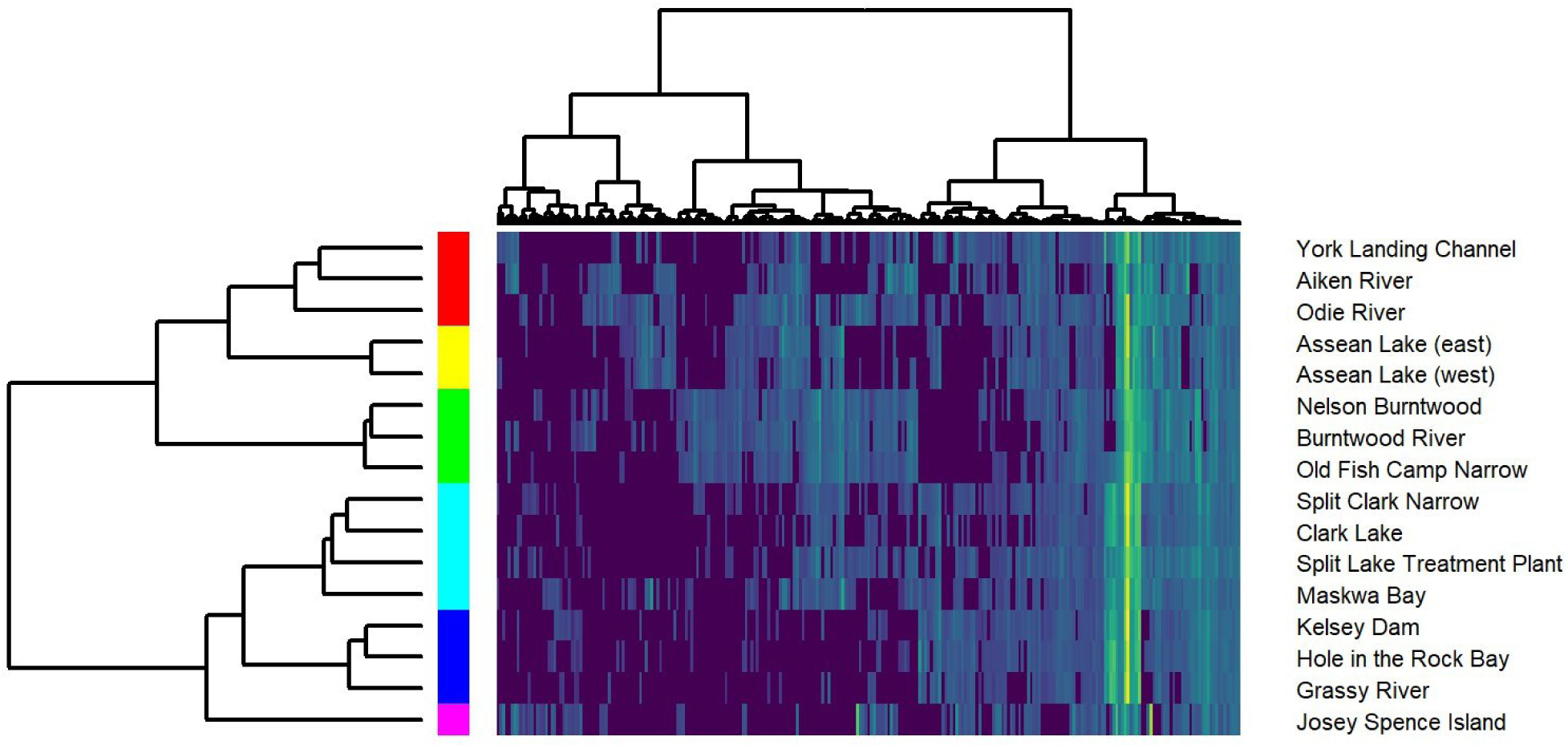
Hierarchical clustering of microbial community composition. Heatmap shows the relative abundance of ASVs (columns) across sites (rows), with darker colors indicating lower relative abundance and brighter colors indicating higher relative abundance. Dendrograms display Ward clustering of Bray–Curtis dissimilarity among taxa (top) and sampling sites (left). The colored bar to the left of the heatmap indicates community clusters.

### 3.5 Microbial community composition

ASVs were taxonomically classified to 23 phyla, 42 classes, 81 orders, 113 families, and 176 genera. Pseudomonadota was the most abundant phylum observed, accounting for 31.6% of the total sequences (Figure 3). The other dominant phyla were Actinomycetota (28.2%), Cyanobacteriota (11.5%), Verrucomicrobiota (9.75%), Chloroflexota (7.30%), and Acidobacteriota (3.82%). Josey Spence Island (JSI) had a distinct profile dominated by Cyanobacteriota (40.2%) and comparatively lower abundance of Verrucomicrobiota (2.9%) and Bacteroidota (1.73%). Nelson Burntwood, Burntwood River, Old Fish Camp Narrow, Split Clark Narrow, Clark Lake, and Split Lake Treatment Plant had a comparatively higher abundance of Acidobacteriota.

**Figure 3.**
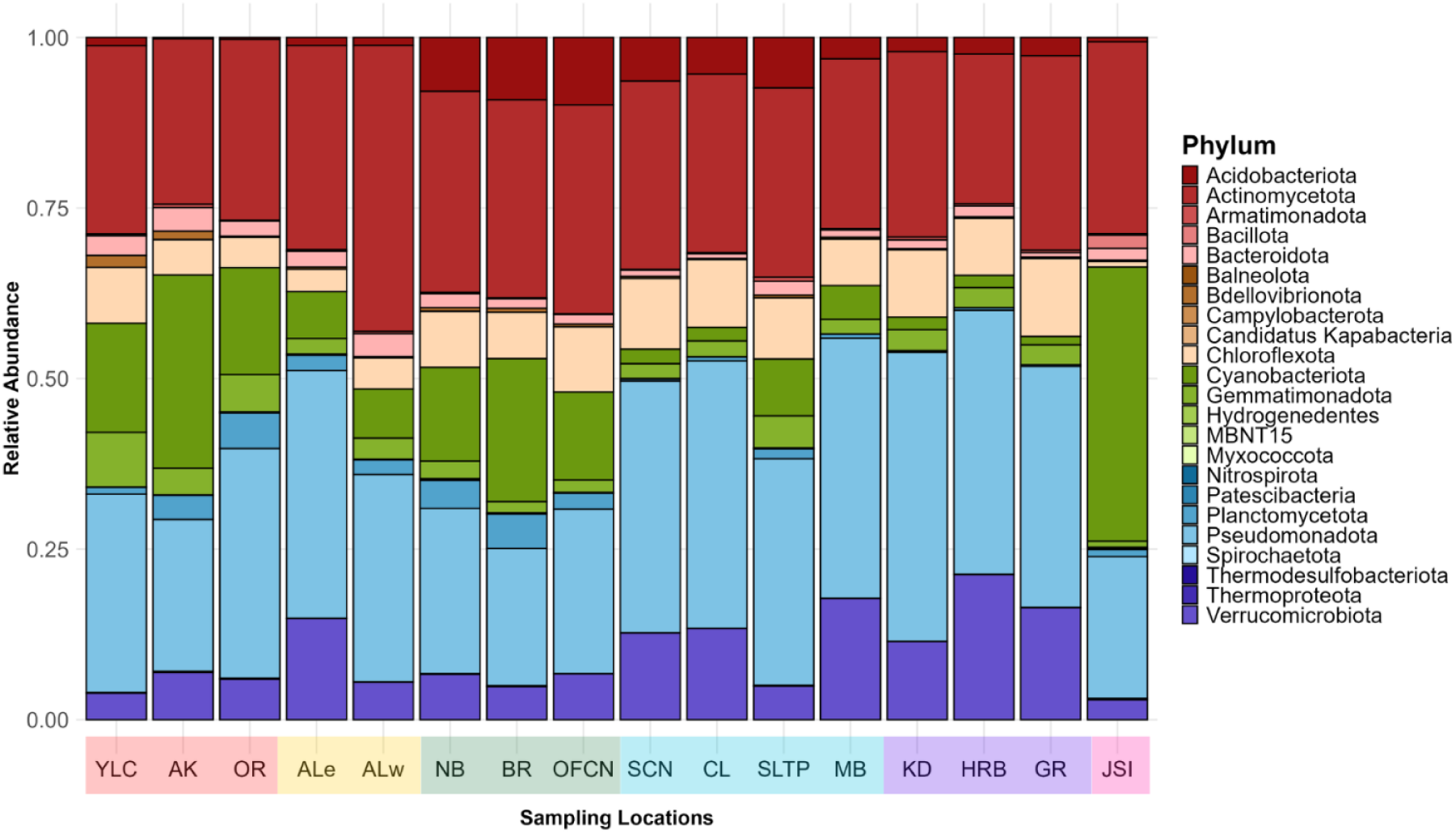
Relative abundance of bacterial phyla across sampling sites. Each bar represents the normalized relative abundance at a given station.

Among all samples, unclassified taxa (incertae sedis) made up the plurality of the relative abundance at the genus level (41.3%); hence clustered together with less abundant genera and were grouped as “other” (Figure 4). The hgcI clade accounted for 16.7% of the community, followed by *Cyanobium* with 10.8%. *Cyanobium* was the most abundant bacterial genus (37.5%) in Josey Spence Island (JSI), suggesting site-specific enrichment, perhaps due to differences in nutrient availability and local hydrodynamics or temporal bloom conditions. Nevertheless, *Cyanobium* abundance was significantly low in Split Clark Narrow, Clark Lake, Kelsey Dam, Hole in the Rock Bay, and Grassy River locations. The “CL500-29 marine group” was found in comparatively high relative abundance at Nelson Burntwood, Burntwood River, Old Fish Camp Narrow, Split Clark Narrow, Clark Lake, and Split Lake Treatment Plant locations. Additionally, the relative abundance of *Polynucleobacter* was comparatively higher in the Aiken River.

**Figure 4.**
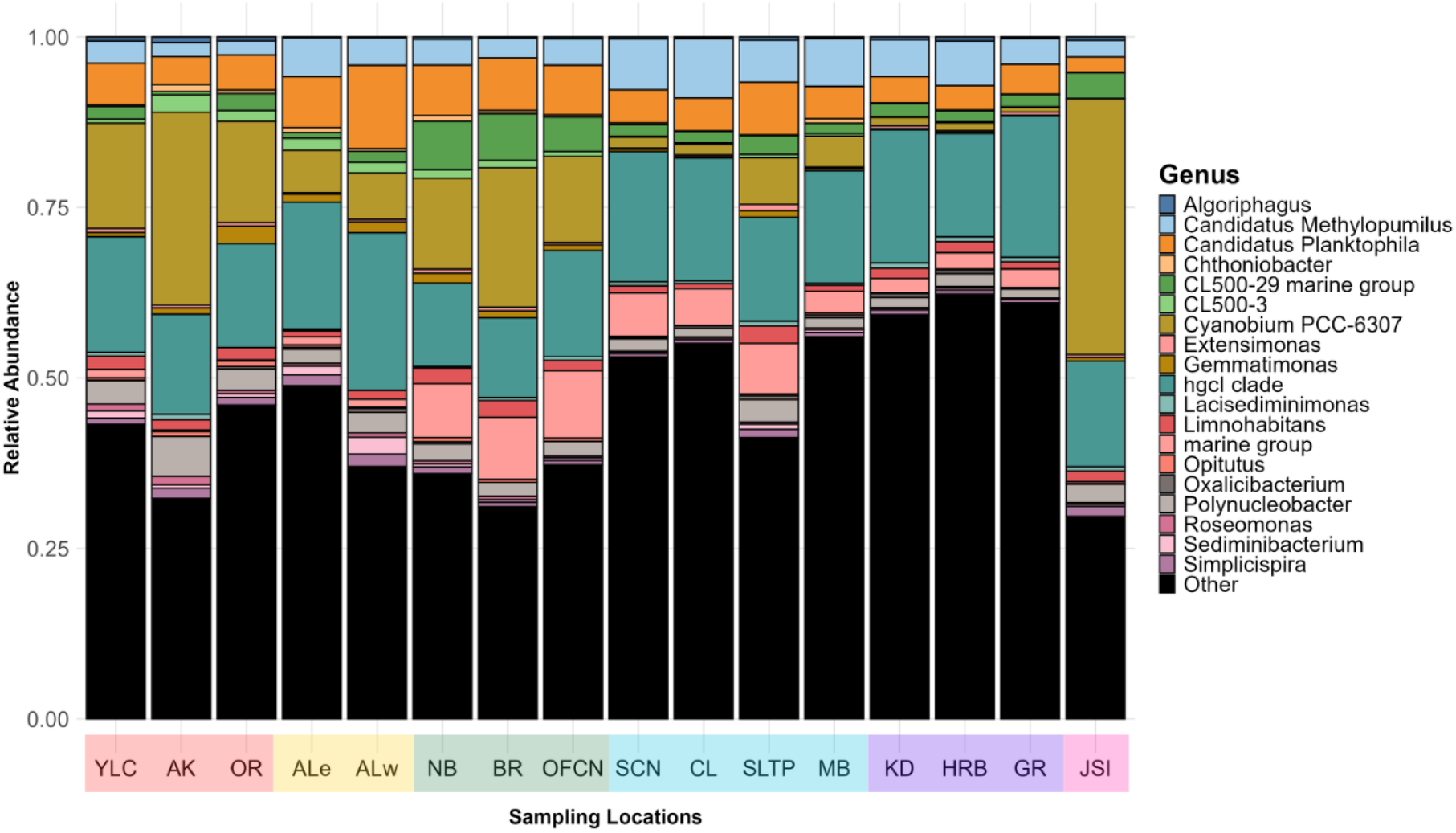
Relative abundance of bacterial genera across sampling sites. The top 20 most abundant genera are shown individually, while all other less abundant genera, including “Incertae Sedis,” are grouped as “Other” (black). Each bar represents the normalized relative abundance at a given station.

### 3.6 Correlation between microbial community and environmental factors

Redundancy analysis (RDA) was performed to evaluate the relationship between bacterial community composition at the ASV level and environmental variables (Figure 5). The first two canonical axes (RDA1 and RDA2) explained the majority of the constrained variation in community composition. Environmental vectors indicated that turbidity and dissolved oxygen were positively associated with RDA1, whereas major ions such as NO_3_*^−^* and Ca^2+^ were aligned primarily along the negative direction of RDA2. However, sample clustering was weak, and most ASVs were distributed near the ordination centroid, suggesting limited structuring of the community by the measured environmental variables. The overall RDA model was not statistically significant based on permutation testing (*F* = 1.63, *p* = 0.057, *N* = 999), indicating that the measured environmental variables explained only a limited proportion of the variation in community composition.

**Figure 5.**
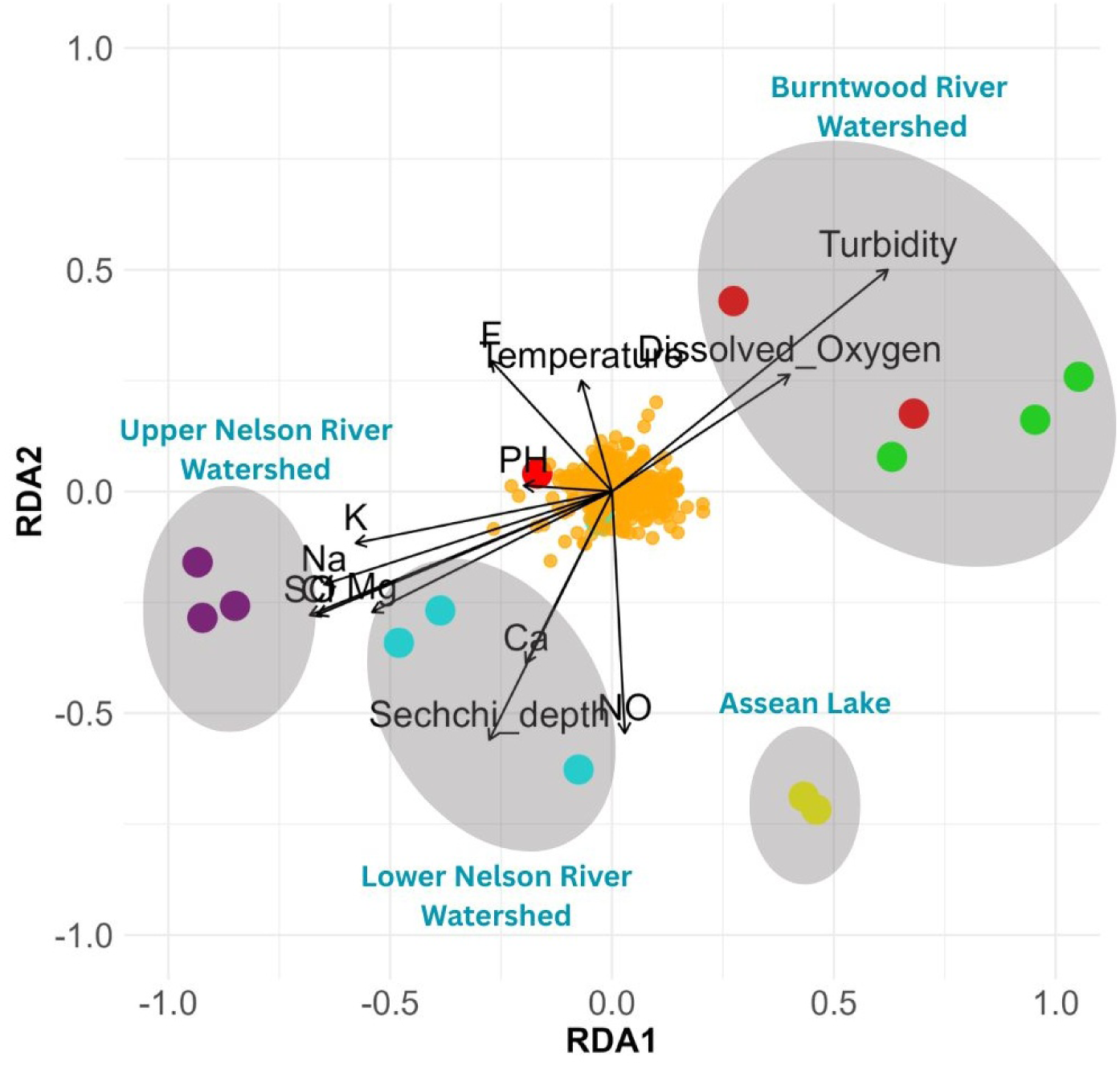
Redundancy analysis (RDA) biplot showing the relationship between ASV community composition (orange dots) and sampling locations (colored dots denote the different community clusters). ASV composition was constrained by environmental parameters, with arrows representing the direction and relative strength of environmental gradients. The watersheds are shown in shaded ellipses. The overall model was not statistically significant based on permutation testing (*F* = 1.63, *p* = 0.057, *N* = 999).

To further assess potential associations between bacterial phyla and environmental variables, Spearman rank correlations were calculated and visualized as a heatmap (Figure S1). Consistent with the RDA results, no statistically significant correlations were detected after Benjamini–Hochberg false discovery rate correction (adjusted *p <* 0.05). Together, these analyses indicate that the measured environmental variables did not exert strong or consistent control on bacterial community composition across samples.

### 3.7 Correlation among microbial phyla

Notably, Pseudomonadota abundance was strongly and negatively correlated with Cyanobacteriota (*R* = *−*0.87, *p* = 0.002), as were the Cyanobacteriota and Verrucomicrobiota (*R* = *−*0.80, *p* = 0.015). Strong positive correlations were observed between Cyanobacteriota and Planctomycetota (*R* = +0.76, *p* = 0.029), Myxococcota and Planctomycetota (*R* = +0.78, *p* = 0.018), and Pseudomonadota and Verrucomicrobiota (*R* = +0.74, *p* = 0.036; Figure 6).

**Figure 6.**
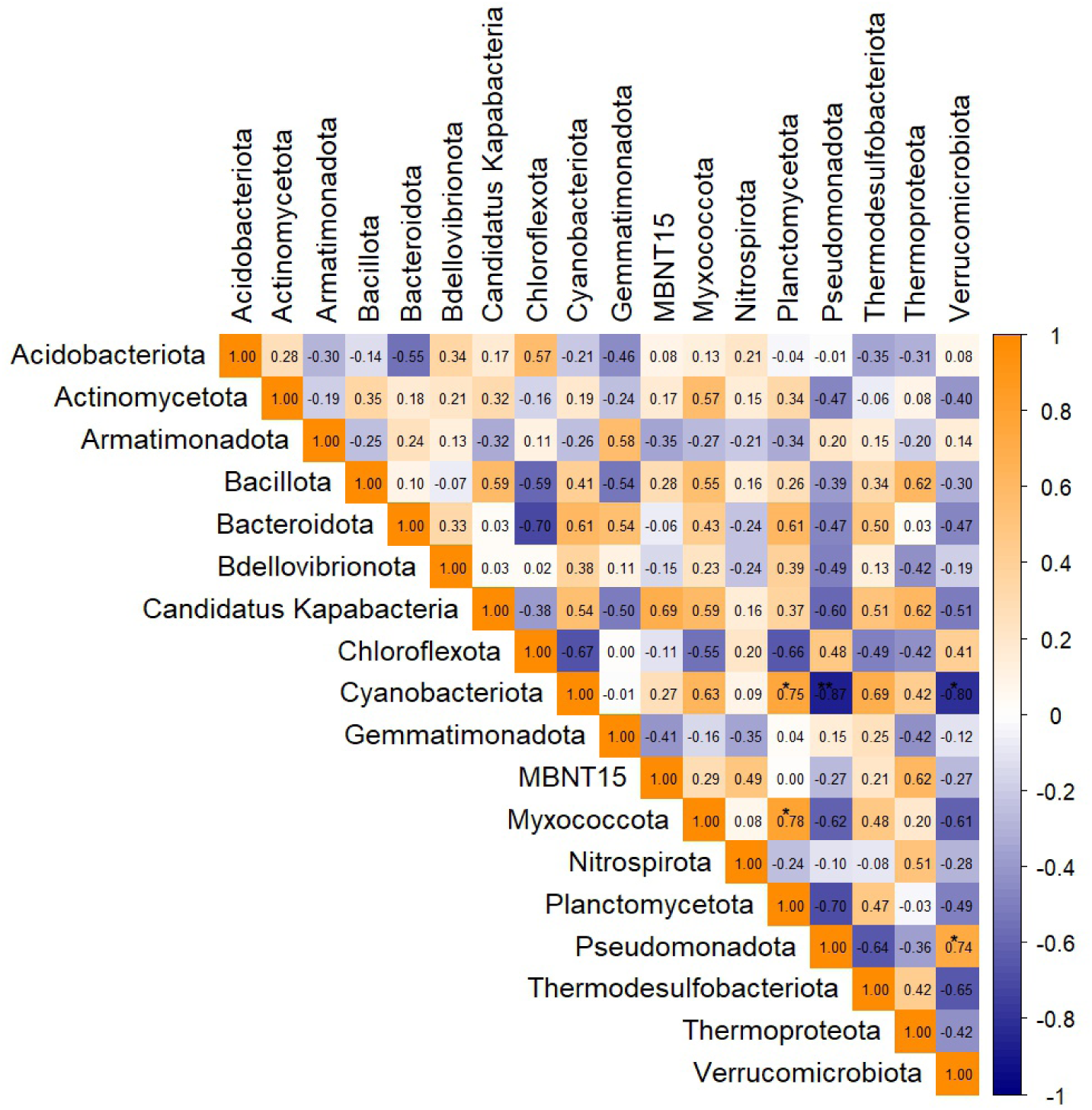
Spearman correlation matrix showing pairwise relationships between bacterial phyla observed in the samples. The color gradient represents the strength and direction of the correlation, with orange indicating positive correlations and blue indicating negative correlations. Values in each cell represent the Spearman correlation coefficient (*ρ*). P-values were adjusted using the Benjamini–Hochberg false discovery rate (FDR) procedure. Asterisks denote significance thresholds (\**p <* 0.05, \*\**p <* 0.01, \*\*\**p <* 0.001)

## 4 Discussion

This study establishes a baseline for microbial community composition in Split Lake, Manitoba, providing a foundation for future investigations across other freshwater lakes in the region. Environmental explanatory variables (including surface water chemistry), dependent on geography, were evaluated as potential drivers of microbial community structure. Freshwater ecosystems host diverse microbial communities that play key roles in nutrient cycling, water quality, and ecosystem functioning. Understanding the environmental structure of these communities is critical in large freshwater lake systems, such as Split Lake, Manitoba. In this study, we used 16S rRNA metabarcoding to characterize microbial community composition across Split Lake in August 2024. We evaluated the relationships between community structure and measured environmental variables using correlation-based statistical approaches. Previous research from Lake Winnipeg, located several hundred kilometers upstream of Split Lake, has shown that bacterial community composition across the lake was relatively stable, with only moderate seasonal variability (Kuzyk et al., 2022). In contrast, microbial communities in Split Lake exhibited pronounced spatial heterogeneity across sampling sites. For example, the Josey Spence Island (JSI) site supported a distinctly different microbial community structure compared to all other sites. This spatial variability likely reflects the stronger influence of localized river inflows, hydrological regulation, and physicochemical gradients within Split Lake, which may override the homogenizing effects observed in larger, more hydrodynamically stable systems such as Lake Winnipeg. JSI exhibited the highest relative abundance of *Cyanobium*, coinciding with very low sulfate concentrations (0.44 ppm), chloride concentrations (0.24 ppm), and moderately elevated turbidity (36.6 NTU). A similar combination of sulfate, chloride, and turbidity was observed at the Odei River site; however, these sites did not cluster together based on the microbial community composition. Given that other measured environmental variables were largely similar between these sites, the contrasting microbial assemblages likely reflect differences in hydrological connectivity, water residence time, or biological interactions rather than direct chemical control, as outlined in previous studies on freshwater lakes’ microbial community dynamics (Kent et al., 2007; Sadeghi et al., 2021, 2024).

Across all sampling sites, Actinobacteria represented the most abundant taxa in Split Lake and its associated inflows. This dominance is consistent with the observations from Lake Winnipeg (Kuzyk et al., 2022), other Canadian freshwater lakes (Luo et al., 2022; Sharma et al., 2009), and all around the world (Allgaier and Grossart, 2006; Newton et al., 2007), where Actinobacteria are frequently among the most prevalent taxa due to their small cell size, high surface area-to-volume ratio, and efficient utilization of low molecular weight dissolved organic carbon (Yaradoddi and Kontro, 2021). Notably, Actinobacteria remained abundant despite pronounced spatial heterogeneity in turbidity, sulfate concentrations, and hydrological influence, suggesting that this group occupies a broadly resilient ecological niche across the Split Lake system. A similar spatial pattern was observed for *Candidatus Planktophila*, a dominant lineage within the Actinobacteriota, which was consistently abundant across all sampling sites. *Candidatus Planktophila* is a ubiquitous freshwater taxon known for its adaptation to oligotrophic conditions, allowing it to persist across a wide range of physicochemical environments (Amadei Martínez et al., 2025).

Hierarchical clustering analysis revealed spatial structuring of microbial communities across Split Lake, identifying six distinct microbial clusters among the 16 sampling sites. These clusters were particularly associated with river inflow regions, suggesting that tributary inputs play a predominant role in shaping microbial community composition within the lake. For example, sites influenced by the upper Nelson River watershed were grouped into different microbial clusters than those associated with the Burntwood River watershed, despite both rivers discharging into Split Lake.

This differentiation likely reflects contrasts in upstream watershed characteristics, major ion and organic matter inputs, and physicochemical conditions associated with each river system. In particular, sulfate concentrations were relatively high in the upper Nelson River watershed (14.46 to 33.32 ppm), compared to the Burntwood River watershed (1.35 to 2.55 ppm), providing a potential geochemical driver for the distinct microbial communities associated with each river system. The increased relative abundance of Verrucomicrobiota and reduced relative abundance in Cyanobacteriota further distinguish these sites from the Burntwood River and are consistent with previous studies on Verrucomicrobiota that describe their metabolic capacity in organic matter degradation, including complex sulfated polysaccharides (SPs) (Sichert et al., 2020; Tan et al., 2025). Spearman correlation revealed several strong negative associations among dominant bacterial phyla, including Cyanobacteriota and Verrucomicrobiota, and Cyanobacteriota and Pseudomonadota. These inverse relationships likely reflect differences in ecological strategies and environmental preferences rather than direct biotic interactions. Many of the negatively associated phyla were also linked to contrasting physicochemical conditions, such as turbidity and dissolved oxygen level, underscoring the relevance of environmental filtering on community structure in Split Lake.

Burntwood River inflows were characterized by higher turbidity levels, and the sampling sites within the lake exhibiting similar turbidity levels to those of the Burntwood River supported microbial assemblages that clustered together and were distinct from communities associated with Nelson River watersheds, Aiken River inflows, and Assean Lake. Elevated turbidity can reduce light penetration and increase particulate matter availability, thereby favoring heterotrophic and particle-associated bacteria while constraining phototrophic taxa (McGowan et al., 2018; Pick and Caron, 1987). These findings indicate that physical drivers such as turbidity act alongside chemical gradients to independently structure microbial communities across Split Lake. The green microbial cluster was characterized by a relatively high abundance of the “CL500-29 marine group” and was primarily associated with Burntwood River inflows and the lower Nelson River watershed, which exhibited elevated turbidity. The CL500-29 clade has been reported in freshwater and brackish systems, where the organic matter is rich (Guo et al., 2023; Wang et al., 2024a,b). The higher turbidity observed in the Burntwood River watershed likely reflects increased suspended particles and organic matter, which may create favorable niches for the CL500-29. In contrast, lower-turbidity inflows and adjacent unregulated waters did not support similar assemblages, underscoring the importance of physical forcing in shaping microbial distributions.

Assean Lake, an adjacent and unregulated freshwater system, lacking direct hydroelectric power influence, exhibited distinct physicochemical and microbial characteristics compared to Split Lake and its inflows. The markedly low turbidity in Assean Lake likely promotes greater light penetration and reduced particle-associated disturbance, conditions that favor free-living, oligotrophic bacterial lineages. The comparatively high relative abundance of the hgcI clade (Actinobacteria) under these conditions is consistent with studies describing this group as adapted to low-nutrient, low-turbidity freshwater environments (Neuenschwander et al., 2018; Yaradoddi and Kontro, 2021).

Prairie lakes, including Lake Winnipeg, are well-documented hotspots for cyanobacterial blooms and elevated microcystin concentrations compared to many other Canadian freshwater systems (McKindles et al., 2019; Onana et al., 2025). In contrast, measured microcystin concentrations in Split Lake remained below maximum acceptable guideline values during the study period. This difference likely reflects the distinct hydrological and physicochemical characteristics of Split Lake, including strong riverine influence, higher turbidity, and reduced water residence times, which can limit bloom development and toxin accumulation. Additionally, although cyanobacterial taxa were present across sites, toxin-producing lineages did not dominate the microbial community, further supporting the observed low microcystin levels.

## 5 Limitations and summary

The present study used only high-throughput 16S rRNA gene sequencing data to explore the microbial community composition in Split Lake. However, an in-depth analysis would be required to understand the community dynamics and functional roles. For instance, these data only cover August 2024; hence, more sampling is required to know how the microbial composition changes between seasons and across years. The sampling primarily focused on the river inflows to Split Lake, with limited data available from the midstream region of the lake. In addition, the potential contribution of particle-associated bacteria cannot be excluded since only surface water samples were collected. Year-round long-term investigations would facilitate identifying the microbial community composition trends and potential ecosystem disruptions, including bloom-forming events, sediment loading, zebra mussel infestation, and contaminant accumulations. Furthermore, metagenomic-level investigations may provide insights into microbial community dynamics, their functional roles, and the underlying genetic mechanisms.

## 6 Conclusion

This study characterized the bacterial community structure of Split Lake, Manitoba, during the summer of 2024 and demonstrated that spatial variability in microbial composition is strongly influenced by environmental and hydrological gradients within the system. Although Actinobacteria, particularly the hgcI clade, and *Cyanobium* dominated most sampling sites, the JSI site exhibited a distinctly different microbial profile, with an unusually high relative abundance of *Cyanobium* compared to all other sites. Hierarchical clustering further revealed six distinct microbial assemblages across Split Lake, highlighting pronounced spatial heterogeneity in community structure. This spatial distribution is likely driven by the unique environmental and hydrological influences in different parts of the lake since Split Lake receives inflows from major rivers, including Burntwood River, Nelson River, Grassy River, and potentially others, each contributing varying concentrations of major ions, sediment loads, and temperature regimes, and shaping the localized environmental conditions for selected microbiota.

Additionally, physical parameters such as flow rate, flow direction, sediment deposition, and light penetration may support niche partitioning among microbial taxa. Therefore, this hydrological complexity, coupled with environmental variables, likely contributes to the spatial diversity of microbiota in Split Lake.

## Data availability

All raw 16S rRNA gene sequence reads generated from this experiment have been deposited at NCBI’s Sequence Read Archive (SRA) under the BioProject PRJNA1416953 (accession numbers SAMN54985744 to SAMN54985759).

## Author contributions

Thamali Vidanage – Conceptualization, Data curation, Formal analysis, Investigation, Methodology, Visualization, Writing – original draft, Writing – review & editing, Project administration

Tataskweyak Environmental Monitoring Agency – Conceptualization, Methodology, Investigation, Resources, Project administration

R. Eric Collins – Conceptualization, Methodology, Resources, Writing – review & editing, Visualization, Validation, Supervision, Funding acquisition, Project administration

## Acknowledgements

We would like to express heartfelt gratitude to the Tataskweyak Cree Nation (TCN) community of the Split Lake #171 and the TCN guardians for the fieldwork support. We extend our sincere gratitude to Dr. Stephane McLachlan and Jeremy Mumford from the Environmental Conservation Lab at the University of Manitoba for their invaluable contribution to this project, and to Debbie Armstrong for performing Ion Chromatography (IC) on the water samples.

## Supplementary material

**Figure S1.**
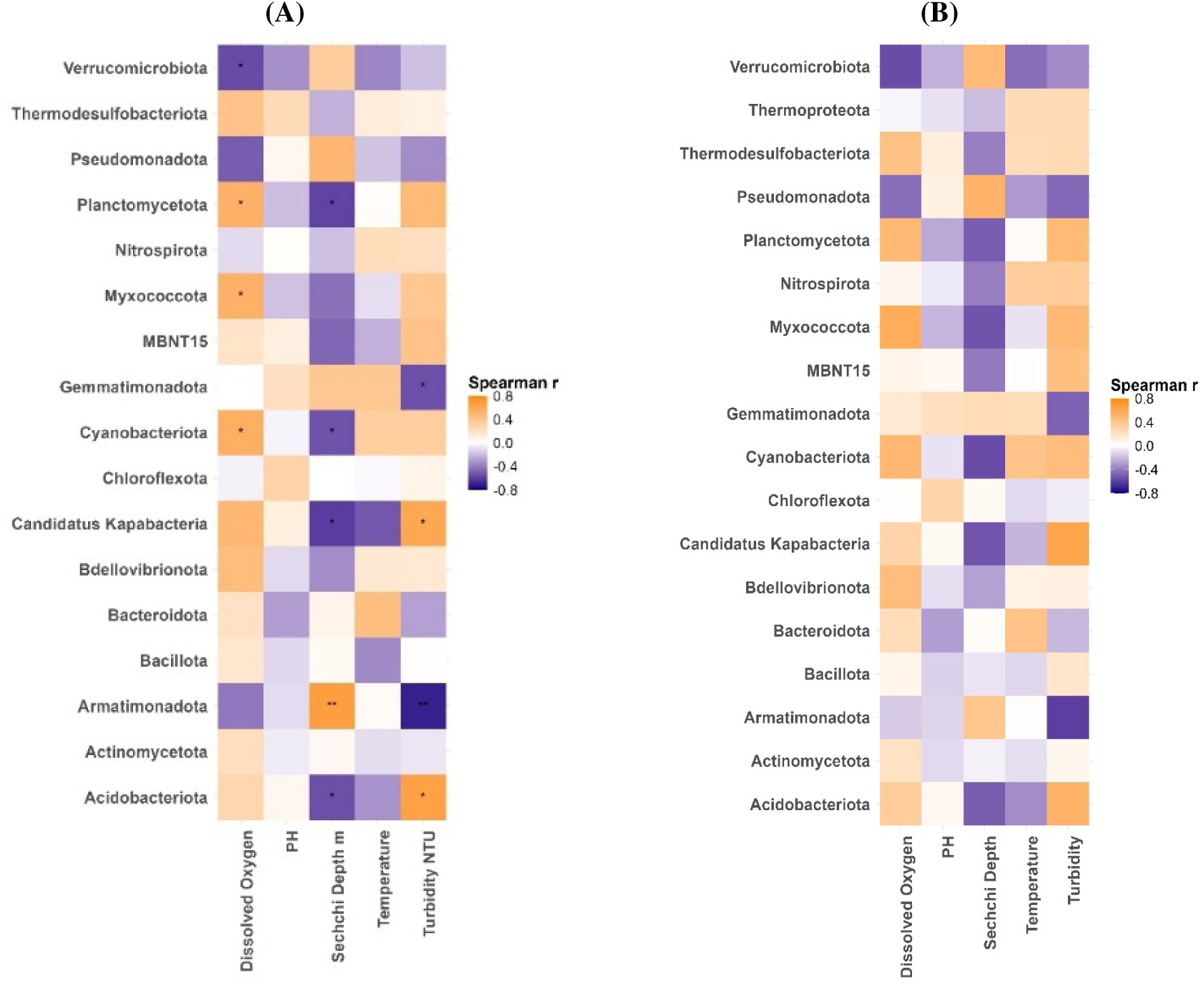
Pairwise Spearman correlations between bacterial phyla and environmental variables. (A) Raw p-values were used to determine statistical significance. (B) P-values were adjusted using the Benjamini–Hochberg false discovery rate (FDR) procedure. Correlation strength is indicated by color intensity (Spearman’s r). Asterisks denote significance thresholds (\**p <* 0.05, \*\**p <* 0.01, \*\*\**p <* 0.001).

**Table S1.** Concentrations of the major cations and anions of the water samples (values are given in ppm). Site ID Mg^2+^ Ca^2+^ Na^+^ K^+^ F*^−^* Cl*^−^* SO_4_^2*−*^ NO_3_*^−^* Cluster color.

| Site ID | Mg<sup>2+</sup> | Ca<sup>2+</sup> | Na<sup>+</sup> | K<sup>+</sup> | F<sup>−</sup> | Cl<sup>−</sup> | SO<sub>4</sub><sup>2−</sup> | NO<sub>3</sub><sup>−</sup> | Cluster color |
|---|---|---|---|---|---|---|---|---|---|
| YLC | 11.40 | 20.31 | 13.48 | 1.96 | 0.11 | 11.73 | 28.65 | 0.34 | Red |
| AK | 6.26 | 18.35 | 4.72 | 0.40 | 0.11 | 3.82 | 9.51 | 0.53 |  |
| OR | 2.56 | 14.77 | 0.74 | <DL | 0.09 | 0.24 | 0.45 | <DL |  |
| ALe | 5.46 | 22.12 | 1.97 | 0.2 | <DL | 0.81 | 1.22 | 0.58 | Yellow |
| ALw | 4.56 | 20.26 | 1.62 | 0.08 | <DL | 0.67 | 1.10 | 0.56 |  |
| NB | 4.38 | 15.93 | 2.96 | 0.66 | 0.11 | 0.93 | 2.55 | <DL | Green |
| BR | 1.58 | 8.11 | 1.02 | <DL | 0.10 | 0.38 | 1.35 | 0.39 |  |
| OFCN | 5.19 | 16.46 | 4.35 | 0.86 | 0.10 | 0.38 | 1.35 | 0.38 |  |
| SCN | 9.38 | 18.88 | 10.61 | 1.69 | 0.11 | 8.97 | 21.93 | 0.45 | Cyan |
| CL | 1.79 | 10.95 | 1.80 | <DL | 0.09 | 1.34 | 4.83 | <DL |  |
| SLTP | 7.79 | 17.62 | 8.10 | 1.34 | 0.11 | 5.98 | 15.06 | 0.36 |  |
| MB | 9.40 | 23.25 | 9.84 | 1.52 | 0.12 | 7.93 | 19.27 | 0.42 |  |
| KD | 12.25 | 20.84 | 15.41 | 2.25 | 0.11 | 13.52 | 33.32 | 0.38 | Purple |
| HRB | 10.74 | 20.60 | 13.12 | 1.84 | 0.11 | 11.39 | 28.66 | 0.37 |  |
| GR | 5.23 | 12.53 | 6.28 | 0.66 | 0.10 | 5.35 | 14.46 | 0.33 |  |
| JSI | 3.79 | 14.70 | 2.84 | 0.62 | 0.09 | 0.24 | 0.44 | <DL | Magenta |

## References

M. Allgaier and H.-P. Grossart. Diversity and seasonal dynamics of Actinobacteria populations in four lakes in Northeastern Germany. Applied and Environmental Microbiology, 72(5):3489–3497, 2006. doi: 10.1128/AEM.72.5.3489-3497.2006.

L. Amadei Martínez, K. Sabbe, S. D’hondt, R. Dasseville, I. Daveloose, T. Verstraete, P. Chaerle, N. Brion, T. Maris, P. Meire, and W. Vyverman. Freshwater discharge and salinity drive taxonomic and functional turnover of microbial communities in a turbid macrotidal estuary. Environmental Microbiology Reports, 17 (4):e70135, 2025. doi: 10.1111/1758-2229.70135.

ASTM. Standard Test Method for Determination of Dissolved Alkali and Alkaline Earth Cations and Ammonium in Water and Wastewater by Ion Chromatography, 2021. ASTM D6919-03.

A. J. Bramburger, C. T. Filstrup, E. D. Reavie, C. S. Sheik, G. D. Haffner, D. C. Depew, and J. A. Downing. Paradox versus paradigm: A disconnect between understanding and management of freshwater cyanobacterial harmful algae blooms. Freshwater Biology, 68(2):191–201, 2023. doi: 10.1111/fwb.14019.

B. J. Callahan, P. J. McMurdie, M. J. Rosen, A. W. Han, A. J. A. Johnson, and S. P. Holmes. DADA2: High-resolution sample inference from Illumina amplicon data. Nature Methods, 13(7):581–583, 2016. doi: 10.1038/nmeth.3869.

Canadian Council of Ministers of the Environment. Canadian Council of Ministers of the Environment, 2007. URL https://ccme.ca/en/resources/water-aquatic-life.

W. W. Carmichael and G. L. Boyer. Health impacts from cyanobacteria harmful algae blooms: Implications for the North American Great Lakes. Harmful Algae, 54:194–212, 2016. doi: 10.1016/j.hal.2016.02.002.

W. Chen, G. Wilkes, I. U. H. Khan, K. D. M. Pintar, J. L. Thomas, C. A. Lévesque, J. T. Chapados, E. Topp, and D. R. Lapen. Aquatic bacterial communities associated with land use and environmental factors in agricultural landscapes using a metabarcoding approach. Frontiers in Microbiology, 9, 2018. doi: 10.3389/fmicb.2018.02301.

A. P. Dufour. Bacterial indicators of recreational water quality. Canadian Journal of Public Health / Revue Canadienne de Santé Publique, 75(1):49–56, 1984. URL https://www.jstor.org/stable/41990233.

Duncan and Williamson. Water Chemistry/ Water Discharge Relationships Within the Churchill River Diversion and Lake Winnipeg Regulation Region, Manitoba, Canada, 1988. URL https://waves-vagues.dfo-mpo.gc.ca/Library/40681609.pdf.

G. Grill, B. Lehner, M. Thieme, B. Geenen, D. Tickner, F. Antonelli, S. Babu, P. Borrelli, L. Cheng, H. Crochetiere, H. Ehalt Macedo, R. Filgueiras, M. Goichot, J. Higgins, Z. Hogan, B. Lip, M. E. McClain, J. Meng, M. Mulligan, C. Nilsson, J. D. Olden, J. J. Opperman, P. Petry, C. Reidy Liermann, L. Sáenz, S. Salinas-Rodríguez, P. Schelle, R. J. P. Schmitt, J. Snider, F. Tan, K. Tockner, P. H. Valdujo, A. van Soesbergen, and C. Zarfl. Mapping the world’s free-flowing rivers. Nature, 569(7755):215–221, 2019. doi: 10.1038/s41586-019-1111-9.

A. D. Gronewold, M. E. Borsuk, R. L. Wolpert, and K. H. Reckhow. An assessment of fecal indicator bacteria-based water quality standards. Environmental Science & Technology, 42(13):4676–4682, 2008. doi: 10.1021/es703144k.

Y. Guo, S. Gu, K. Wu, A. J. Tanentzap, J. Yu, X. Liu, Q. Li, P. He, D. Qiu, Y. Deng, P. Wang, Z. Wu, and Q. Zhou. Temperature-mediated microbial carbon utilization in China’s lakes. Global Change Biology, 29 (17):5044–5061, 2023. doi: 10.1111/gcb.16840.

S. Hermans, A. Gautam, G. D. Lewis, M. Neale, H. L. Buckley, B. S. Case, and G. Lear. Exploring freshwater stream bacterial communities as indicators of land use intensity. Environmental Microbiome, 19(1):45, 2024. doi: 10.1186/s40793-024-00588-z.

M. M. M. Islam, S. Shafi, S. A. Bandh, and N. Shameem. Chapter 3—Impact of environmental changes and human activities on bacterial diversity of lakes. In S. A. Bandh, S. Shafi, and N. Shameem, editors, Freshwater Microbiology, pages 105–136. Academic Press, 2019. doi: 10.1016/B978-0-12-817495-1.00003-7.

P. Joshi. Microbial diversity of aquatic ecosystem and its industrial potential. Journal of Bacteriology & Mycology: Open Access, 3(1), 2016. doi: 10.15406/jbmoa.2016.03.00048.

A. D. Kent, A. C. Yannarell, J. A. Rusak, E. W. Triplett, and K. D. McMahon. Synchrony in aquatic microbial community dynamics. The ISME Journal, 1(1):38–47, 2007. doi: 10.1038/ismej.2007.6.

S. B. Kuzyk, X. Ma, and V. Yurkov. Seasonal dynamics of Lake Winnipeg’s microbial communities reveal aerobic anoxygenic phototrophic populations coincide with sunlight availability. Microorganisms, 10(9): 1690, 2022. doi: 10.3390/microorganisms10091690.

J. Luo, H. Zeng, Q. Zhou, X. Hu, Q. Qu, S. Ouyang, and Y. Wang. Anthropogenic impacts on the biodiversity and anti-interference ability of microbial communities in lakes. Science of The Total Environment, 820: 153264, 2022. doi: 10.1016/j.scitotenv.2022.153264.

Manitoba Hydro. Regional Cumulative Effects Assessment. Technical report, Manitoba Hydro, dec 2015. URL https://www.hydro.mb.ca/docs/regulatory_affairs/pdf/rcea/rcea_phase2_part_v_water_part_1.pdf.

M. Martin. Cutadapt removes adapter sequences from high-throughput sequencing reads. EMBnet.Journal, 17(1):10–12, 2011. doi: 10.14806/ej.17.1.200.

M. Mašín, J. Jezbera, J. Nedoma, V. Straškrabová, J. Hejzlar, and K. Šimek. Changes in bacterial community composition and microbial activities along the longitudinal axis of two canyon-shaped reservoirs with different inflow loading. Hydrobiologia, 504(1–3):99–113, 2003. doi: 10.1023/B:HYDR.0000008512.04563.0b.

S. McGowan, N. J. Anderson, M. E. Edwards, E. Hopla, V. Jones, P. G. Langdon, A. Law, N. Solovieva, S. Turner, M. van Hardenbroek, E. J. Whiteford, and E. Wiik. Vegetation transitions drive the autotrophy– heterotrophy balance in Arctic lakes. Limnology and Oceanography Letters, 3(3):246–255, 2018. doi: 10.1002/lol2.10086.

K. M. McKindles, P. V. Zimba, A. S. Chiu, S. B. Watson, D. B. Gutierrez, J. Westrick, H. Kling, and T. W. Davis. A multiplex analysis of potentially toxic Cyanobacteria in Lake Winnipeg during the 2013 bloom season. Toxins, 11(10):587, 2019. doi: 10.3390/toxins11100587.

J. S. Metcalf and G. A. Codd. Cyanotoxins. In B. A. Whitton, editor, Ecology of Cyanobacteria II: Their Diversity in Space and Time, pages 651–675. Springer Netherlands, 2012. doi: 10.1007/978-94-007-3855-3_24.

J. Méndez, A. Audicana, M. Cancer, A. Isern, J. Llaneza, B. Moreno, M. Navarro, M. L. Tarancón, F. Valero, F. Ribas, J. Jofre, and F. Lucena. Assessment of drinking water quality using indicator bacteria and bacteriophages. Journal of Water and Health, 2(3):201–214, 2004. doi: 10.2166/wh.2004.0018.

S. M. Neuenschwander, R. Ghai, J. Pernthaler, and M. M. Salcher. Microdiversification in genome-streamlined ubiquitous freshwater Actinobacteria. The ISME Journal, 12(1):185–198, 2018. doi: 10.1038/ismej.2017.156.

R. J. Newton, S. E. Jones, M. R. Helmus, and K. D. McMahon. Phylogenetic ecology of the freshwater Actinobacteria acI lineage. Applied and Environmental Microbiology, 73(22):7169–7176, 2007. doi: 10.1128/AEM.00794-07.

R. J. Newton, S. E. Jones, A. Eiler, K. D. McMahon, and S. Bertilsson. A guide to the natural history of freshwater lake bacteria. Microbiology and Molecular Biology Reviews, 75(1):14–49, 2011. doi: 10.1128/mmbr.00028-10.

C. Nilsson and K. Berggren. Alterations of riparian ecosystems caused by river regulation: Dam operations have caused global-scale ecological changes in riparian ecosystems. how to protect river environments and human needs of rivers remains one of the most important questions of our time. BioScience, 50(9): 783–792, 2000. doi: 10.1641/0006-3568(2000)050[0783:AORECB]2.0.CO;2.

P. Nowicka-Krawczyk, J. Zelazna Wieczorek, I. Skrobek, M. Ziułkiewicz, M. Adamski, A. Kaminski, and P. Z mudzki. Persistent Cyanobacteria blooms in artificial water bodies—An effect of environmental conditions or the result of anthropogenic change. International Journal of Environmental Research and Public Health, 19(12):6990, 2022. doi: 10.3390/ijerph19126990.

J. Oksanen, G. L. Simpson, F. G. Blanchet, R. Kindt, P. Legendre, P. R. Minchin, R. B. O’Hara, P. Solymos, M. H. H. Stevens, E. Szoecs, H. Wagner, M. Barbour, M. Bedward, B. Bolker, D. Borcard, T. Borman, G. Carvalho, M. Chirico, M. D. Caceres, et al. vegan: Community Ecology Package, 2025. URL https://cran.r-project.org/web/packages/vegan/index.html. Version 2.7-1 [Computer software].

V. E. Onana, B. E. Beisner, and D. A. Walsh. Water quality and land use shape bacterial communities across 621 Canadian lakes. Environmental Microbiology, 27(1):e70037, 2025. doi: 10.1111/1462-2920.70037.

J. D. Pfaff. Determination of inorganic anions by ion chromatography. In Methods for the Determination of Metals in Environmental Samples, pages 388–417. Elsevier, 1996. doi: 10.1016/B978-0-8155-1398-8.50022-7.

F. R. Pick and D. A. Caron. Picoplankton and nanoplankton biomass in Lake Ontario: Relative contribution of phototrophic and heterotrophic communities. Canadian Journal of Fisheries and Aquatic Sciences, 44 (12):2164–2172, 1987. doi: 10.1139/f87-265.

C. Quast, E. Pruesse, P. Yilmaz, J. Gerken, T. Schweer, P. Yarza, J. Peplies, and F. O. Glöckner. The SILVA ribosomal RNA gene database project: Improved data processing and web-based tools. Nucleic Acids Research, 41(D1):D590–D596, 2013. doi: 10.1093/nar/gks1219.

J. Sadeghi, S. R. Chaganti, A. H. Shahraki, and D. D. Heath. Microbial community and abiotic effects on aquatic bacterial communities in north temperate lakes. Science of The Total Environment, 781:146771, 2021. doi: 10.1016/j.scitotenv.2021.146771.

J. Sadeghi, C. J. Venney, S. Wright, J. Watkins, D. Manning, E. Bai, C. Frank, and D. D. Heath. Aquatic bacterial community connectivity: The effect of hydrological flow on community diversity and composition. Environments, 11(5):90, 2024. doi: 10.3390/environments11050090.

D. W. Schindler. Recent advances in the understanding and management of eutrophication. Limnology and Oceanography, 51(1part2):356–363, 2006. doi: 10.4319/lo.2006.51.1_part_2.0356.

A. K. Sharma, K. Sommerfeld, G. S. Bullerjahn, A. R. Matteson, S. W. Wilhelm, J. Jezbera, U. Brandt, W. F. Doolittle, and M. W. Hahn. Actinorhodopsin genes discovered in diverse freshwater habitats and among cultivated freshwater Actinobacteria. The ISME Journal, 3(6):726–737, 2009. doi: 10.1038/ismej.2009.13.

A. Sichert, C. H. Corzett, M. S. Schechter, F. Unfried, S. Markert, D. Becher, A. Fernandez-Guerra, M. Liebeke, T. Schweder, M. F. Polz, and J.-H. Hehemann. Verrucomicrobia use hundreds of enzymes to digest the algal polysaccharide fucoidan. Nature Microbiology, 5(8):1026–1039, 2020. doi: 10.1038/s41564-020-0720-2.

X.-Y. Tan, X.-J. Liu, D.-C. Lu, Y.-Q. Ye, X.-Y. Liu, F. Yu, H. Yang, F. Li, Z.-J. Du, and M.-Q. Ye. Insights into the physiological and metabolic features of *thalassobacterium*, a novel genus of Verrucomicrobiota with the potential to drive the carbon cycle. mBio, 16(4):e00305–25, 2025. doi: 10.1128/mbio.00305-25.

N. Thajuddin and D. Dhanasekaran. Algae: Organisms for Imminent Biotechnology. BoD – Books on Demand, 2016.

B. Wang, K. Hu, C. Li, Y. Zhang, C. Hu, Z. Liu, J. Ding, L. Chen, W. Zhang, J. Fang, and H. Zhang. Geographic distribution of bacterial communities of inland waters in China. Environmental Research, 249: 118337, 2024a. doi: 10.1016/j.envres.2024.118337.

Q. Wang, G. Jiang, Z. Sun, Y. Liang, F. Liu, and J. Shi. Water quality and microecosystem of water tanks in karst mountainous area, Southwest China. Environmental Science and Pollution Research, 31(9): 12948–12965, 2024b. doi: 10.1007/s11356-024-31959-1.

X. Wen, F. Chen, Y. Lin, H. Zhu, F. Yuan, D. Kuang, Z. Jia, and Z. Yuan. Microbial indicators and their use for monitoring drinking water quality—a review. Sustainability, 12(6):2249, 2020. doi: 10.3390/su12062249.

A. P. Wiens. Churchill river diversion: Effects on benthic invertebrates in lakes along the lower Churchill and the diversion route. Canadian technical report of fisheries and aquatic sciences No. 2001. Technical Report MIC-95-03854/XAB; SSC-FS-97-6/2001E, Department of Fisheries and Oceans, Winnipeg, MB (Canada). Central and Arctic Region, 1994. URL https://www.osti.gov/etdeweb/biblio/117953.

J. S. Yaradoddi and M. H. Kontro. Actinobacteria: Basic adaptation to harsh environments. In J. S. Yaradoddi, M. H. Kontro, and S. V. Ganachari, editors, Actinobacteria: Ecology, Diversity, Classification and Extensive Applications, pages 69–88. Springer Nature, 2021. doi: 10.1007/978-981-16-3353-9_5.

